# Development of a high-throughput barrier function assay using primary human gut organoids

**DOI:** 10.1101/2022.12.16.520780

**Authors:** Charles W. Wright, Naomi Li, Lynsey Shaffer, Armetta Hill, Nicolas Boyer, Stephen Alves, Sriram Venkat, Kaustav Biswas, Linda A. Lieberman, Sina Mohammadi

## Abstract

Disruptions in the gut epithelial barrier can lead to the development of chronic indications such as inflammatory bowel disease (IBD). Historically, barrier function has been assessed in cancer cell lines, which do not contain all human intestinal cell types, leading to poor translatability. To bridge this gap, we adapted human primary gut organoids grown as monolayers to assess barrier function. In this work we describe and characterize a novel 96-well human gut organoid-derived monolayer system that enables medium/high-throughput quantitative assessment of candidate therapeutics. Normal human intestine differentiation patterns and barrier function were characterized and confirmed to recapitulate key aspects of *in vivo* biology. Next, cellular response to TNF-α (a central driver of IBD) was determined using a diverse cadre of readouts.These outputs included transcription factor phosphorylation, target gene expression, cytokine/chemokine production, and barrier function. Additionally, we showed that TNF-α pathway antagonists rescued damage caused by TNF-α in a dose-dependent manner, indicating that this system is suitable for quantitative assessment of barrier modulating factors. Taken together, we have established a robust primary cell-based 96-well system capable of interrogating questions around mucosal response. This system is well suited to provide pivotal functional data to support translational target and drug discovery efforts.

## Introduction

Inflammatory bowel disease (IBD) is a collection of disorders that present as inflammation of the intestinal mucosa ^1,2^. A hallmark feature of IBD is the loss of epithelial barrier function ^3-5^. The gut epithelium plays two roles in homeostasis: while the epithelium prevents the leakage of luminal content into serosal spaces, it also serves as a conduit, relaying signals to immune cells in the lamina propria ^5^. In IBD the loss of barrier function leads to dysregulated exposure of immune cells to gut luminal content. Additionally, cytokines and chemokines produced by the epithelium (e.g. IL-8, MCPs, IL-1β) exacerbate mucosal inflammation. A key driver in IBD is TNF-α. This cytokine, produced by macrophages, dendritic cells, and T cells plays a pleiotropic role in disease. Importantly, TNF-α causes pronounced barrier dysfunction ^6,7^. Because of the central role for TNF-α, TNF-α-neutralizing therapeutics have been used for decades to treat IBD. However, while initially anti-TNF-α therapeutics are effective, subsets of IBD patients stop responding to these therapies, highlighting the need to develop a better understanding of the role for TNF-α in IBD. The development of high-throughput, translatable *in vitro* systems that recapitulate TNF-α signaling would be pivotal in direct investigation of physiologically relevant biology and enable development of more effective therapeutics.

Current high-throughput gut barrier function models are based on human adenocarcinoma cell lines Caco-2 and HT-29 ^8-10^. A considerable deficit of these models is the lack of cell type complexity found in the human intestine. Four predominant cell types are found in the human intestine: absorptive enterocytes, secretory goblet, enteroendocrine, and Paneth cells ^11^. These cell types carry out the functions of the intestine, thus are important constituents of *in vitro* barrier function cell-based assays. As cell line-based systems do not contain multiple cell types, gut organoid-based systems have been adapted to interrogate barrier function.

Gut organoids are complex cultures that contain most of the cell types found in the human intestine ^12,13^. These cultures have been used to recapitulate important aspects of human disease *in vitro* ^14-16^. An important challenge with organoid cultures is access to both apical and basolateral surfaces of the epithelium. The basolateral surface faces the culture media in organoids grown in 3D, thereby limiting access to the apical surface. Additionally, embedded growth in hydrogels (such as Matrigel), may limit access of treatment compounds or biologics to the basolateral surface. These challenges highlight the need for developing robust systems in which access to both apical and basolateral surface is not hindered by polarity or hydrogels.

Human gut organoid-derived cultures have been adapted to 2D monolayer cultures and used to model a range of biology from infectious diseases to barrier function defects ^17-21^. Developed models have been shown to recapitulate key aspects of biology and disease in relatively low-throughput formats (24- and 12-well systems). While these systems add value for modeling human gut biology, the large format hinders quantitative assessment of candidate therapeutics.

In this work, we describe the development and characterization of a robust 96-well human gut organoid-derived monolayer system. We show the implementation of multiple readouts in this system including quantitative measures of transcription factor phosphorylation, gene expression, cytokine production, and barrier function. We focus on TNF-α-mediated signaling events and show pathway modulation by quantifying the effects of TNF-α inhibitors using multiple assays. This miniaturized system enables high-throughput testing of candidate therapeutics in a complex and physiologically relevant culture system that recapitulates key aspects of the human intestine. These include barrier function, polarization, and production of inflammatory cytokines.

## Results

### Establishing and characterizing organoid-derived monolayer cultures

Human adult stem cell (ASC) gut organoid cultures were established and maintained as previously described ^22,23^ (Figure 1A). Organoid-derived monolayer cultures were established by adapting a previously published protocol ^24^ to human ASC organoids. Notably, 96-well transwell cultures were used to facilitate implementation of automation, thereby increasing throughput, and enabling dose-curve experiments. Importantly, commercially available reagents and robots were used to facilitate ease of implementation in any lab setting. Briefly, 3D organoid cultures were dissociated to single cells and plated onto 96-well transwells (Figure 1A). Cultures were allowed to differentiate by withdrawing Wnt pathway agonists from culture medium (see Methods for details). Monolayer cultures were subsequently characterized for barrier function and differentiation. Barrier function was quantified by measuring transepithelial electrical resistance (TEER), a non-invasive measure of barrier integrity that allowed for longitudinal measurements on the same transwells. This method showed that cultures typically reached a plateau of barrier tightness ∼8 days after seeding (Figure 1B). Importantly, high TEER readings were maintained, enabling multi-day experiments.

**Figure 1:**
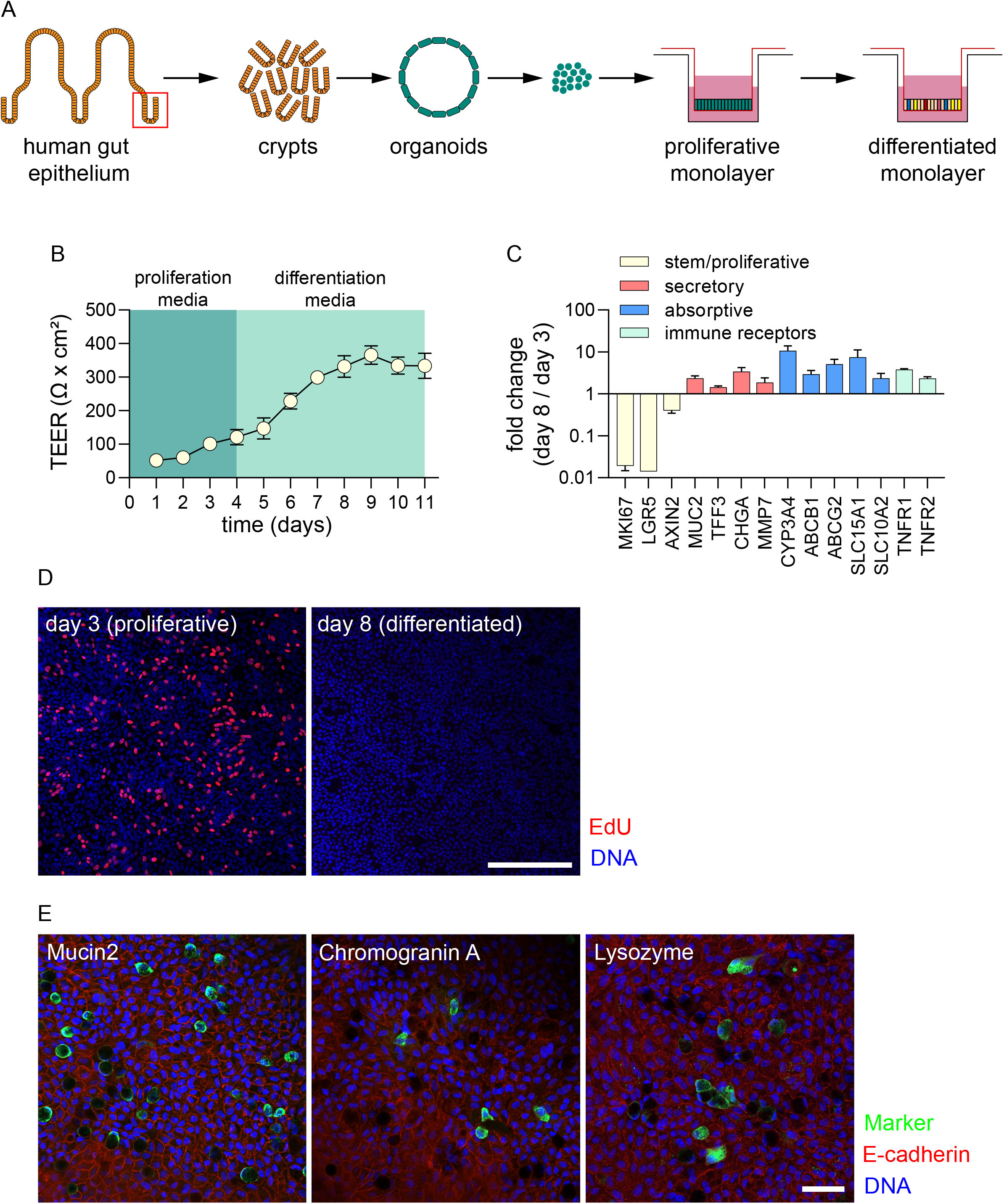
Establishment and characterization of organoid-derived monolayer cultures (A) Schematic of 3D organoid generation and adaptation into 2D organoid-derived monolayers. Stem cell enriched, 3D organoids were generated from crypts isolated from healthy human intestinal epithelial tissue. These organoids were broken into single cells and seeded onto semi-permeable transwells to generate 2D cultures. (B) TEER measurements over a proliferation and differentiation time course. Cultures were grown in proliferation medium for 4 days before switching to differentiation medium for the following 7 days. Barrier function was measured in raw resistance values and normalized by surface area to obtain TEER values. Measurements were taken at approximately 24 hour intervals after seeding for the duration of this culture. Each data point is an average of 8 biological replicates. (C) Gene expression analysis for markers of stem/proliferative (yellow), secretory (red), and absorptive (blue) cells, as well as TNF receptors (green). Values are reported as fold change in differentiated cultures compared to proliferative cultures. Experiment was performed with 3 cell culture replicates. (D) Staining for S-phase cells using EdU assay for proliferative (day 3) and differentiated (day 8 cultures). Pseudocoloring is as follows: blue, DNA; red, proliferative/S-phase cells. Scale bar = 200 µm. (E) Differentiated organoid-derived monolayers were fixed and stained for makers of secretory cell types. Pseudocoloring is as follows: blue, DNA; green, lysozyme, mucin 2, or chromogranin A; red, E-cadherin. Scale bar = 50 µm.

The differentiation status of cultures was determined through assessing presence of functional cell marker through gene expression analysis and immunofluorescence microscopy. Gene expression analysis showed that markers of stem and proliferative cells (LGR5, MKI67, and AXIN2) were downregulated as cultures differentiated (Figure 1C). Importantly, the expression of secretory and absorptive cell markers (secretory: MUC2, TFF3, CHGA, MMP7; absorptive: CYP3A4, ABCB1, ABCG2, SLC15A1, SLC10A2) were upregulated in differentiated cultures, indicating the presence of functional cells (Figure 1C). Concomitantly, differentiated cultures showed a marked reduction in S-phase cells (Figure 1D), indicating the presence of terminally differentiated cells. Furthermore, staining for functional cell markers using immunofluorescence microscopy confirmed the presence of goblet cells, enteroendocrine cells, and Paneth cells (Figure 1E). Taken together, these results indicate that monolayer cultures differentiate to form the expected intestinal cell types while maintaining barrier function.

### Developing a TNF-*α*-based barrier damage system

As TNF-α is a key driver of barrier dysfunction in IBD ^6,7^, barrier damage by TNF was assessed using ASC organoid-derived monolayer cultures. Importantly, gene expression analysis showed that TNF receptors were expressed in differentiated cultures (Figure 1C). Quantification of barrier over time showed that the addition of TNF-α to culture media resulted in a gradual reduction in TEER (Figure 2A). At 24 hours post TNF-α addition, little loss in TEER was observed. However, at 48- and 72-hours post-treatment, a marked reduction in TEER was recorded. Next, the effect of a dose series of TNF-α upon barrier function (TEER) was quantified. This experiment generated a dose-dependent reduction in TEER, measuring a half maximal effective concentration (EC50) of 7 ng/mL (Figure 2B). These results demonstrate that this ASC organoid-derived monolayer system is responsive to TNF-α and could be used to evaluate TNF-α pathway modulators.

**Figure 2:**
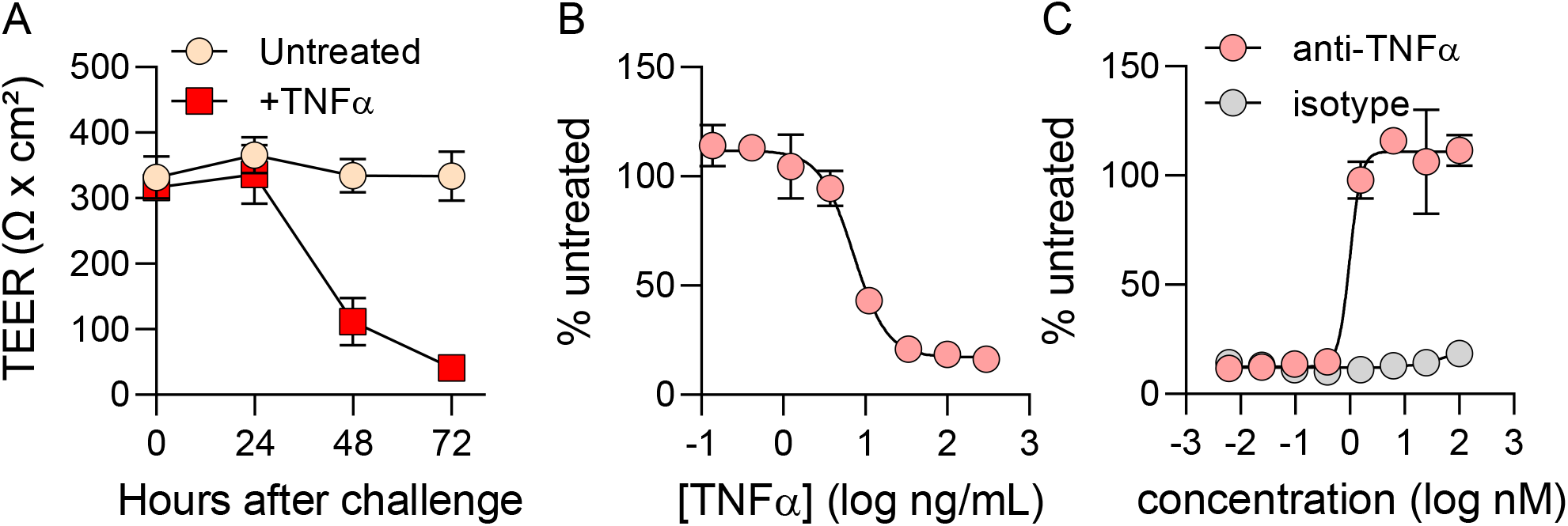
Developing a TNF-α-based barrier damage system (A) Barrier function was quantified in untreated monolayers (yellow) or monolayers challenged with 40 ng/mL TNF-α (red). TEER measurements were taken immediately before challenge and then in 24-hour intervals. Each data point represents the average of 8 cell culture replicates. (B) Barrier disruption following 72-hour challenge with an 8-point dose curve with TNF-α. Values are reported as %untreated, which was obtained by averaging 5 cell culture replicates for each concentration of TNF-α and normalizing to the average TEER value of untreated controls (4 cell culture replicates). Curve fit and EC50 values were generated using a nonlinear regression model in GraphPad Prism. (C) Evaluation of anti-TNF-α antibody for the prevention of barrier damage. Monolayers were pre-treated for 1 hour with 8-point dose curves of either anti-TNF-α neutralizing antibody or an isotype-matched control before challenge with 40 ng/mL TNF-α. TEER measurements were taken at 72-hours post-challenge and plotted in a similar manner to Figure 2B.

The effect of TNF-α pathway modulators upon barrier function was quantified next. TNF-α-targeting biologics have revolutionized treatment for IBD over the last two decades ^25,26^ so the effect of an anti-TNF-α monoclonal antibody was assessed. Addition an anti-TNF-α antibody to monolayer cultures treated with TNF-α efficiently rescued barrier damage in a dose-dependent manner (Figure 2C). The EC50 for the anti-TNF-α antibody was measured to be 1 nM, indicating robust inhibition of TNF-α-mediated damage. Importantly, an isotype antibody used at the same concentrations failed to rescue barrier damage, indicating that this phenotype is specific to the anti-TNF-α antibody activity. These results indicate that our organoid-derived monolayer system could be deployed to quantitatively assess the effectiveness of TNF-α pathway modulators.

### Apical vs basolateral TNF-*α* signaling

Next, we investigated the polarity of TNF-α signaling. Our monolayer system allows for access to both apical and basolateral surfaces of the epithelium, thereby enabling us to parse differences in the location from which signaling is initiated. Monolayer cultures were challenged with TNF-α (as described in Figure 2), however here TNF-α was added either to the apical or basal chamber. Barrier integrity was then quantified using TEER. While apical challenge with TNF-α did very little damage to the barrier, basolateral challenge reduced TEER in a dose-dependent manner (Figure 3A). The EC50 for basolateral TNF-α challenge was measured to be 4 ng/mL, a figure very similar to that observed with TNF-α challenge on both sides of the epithelium (7 ng/mL, Figure 2B). This observation indicates that most of TNF-α signaling is initiated in the basolateral compartment. These results were supported by gene expression analysis. The expression of well characterized TNF-α-responsive genes IL1B and TNF were quantified using quantitative PCR, showing that gene expression was stimulated only when TNF-α was added in the basal chamber of transwells (Figure 3B).

**Figure 3:**
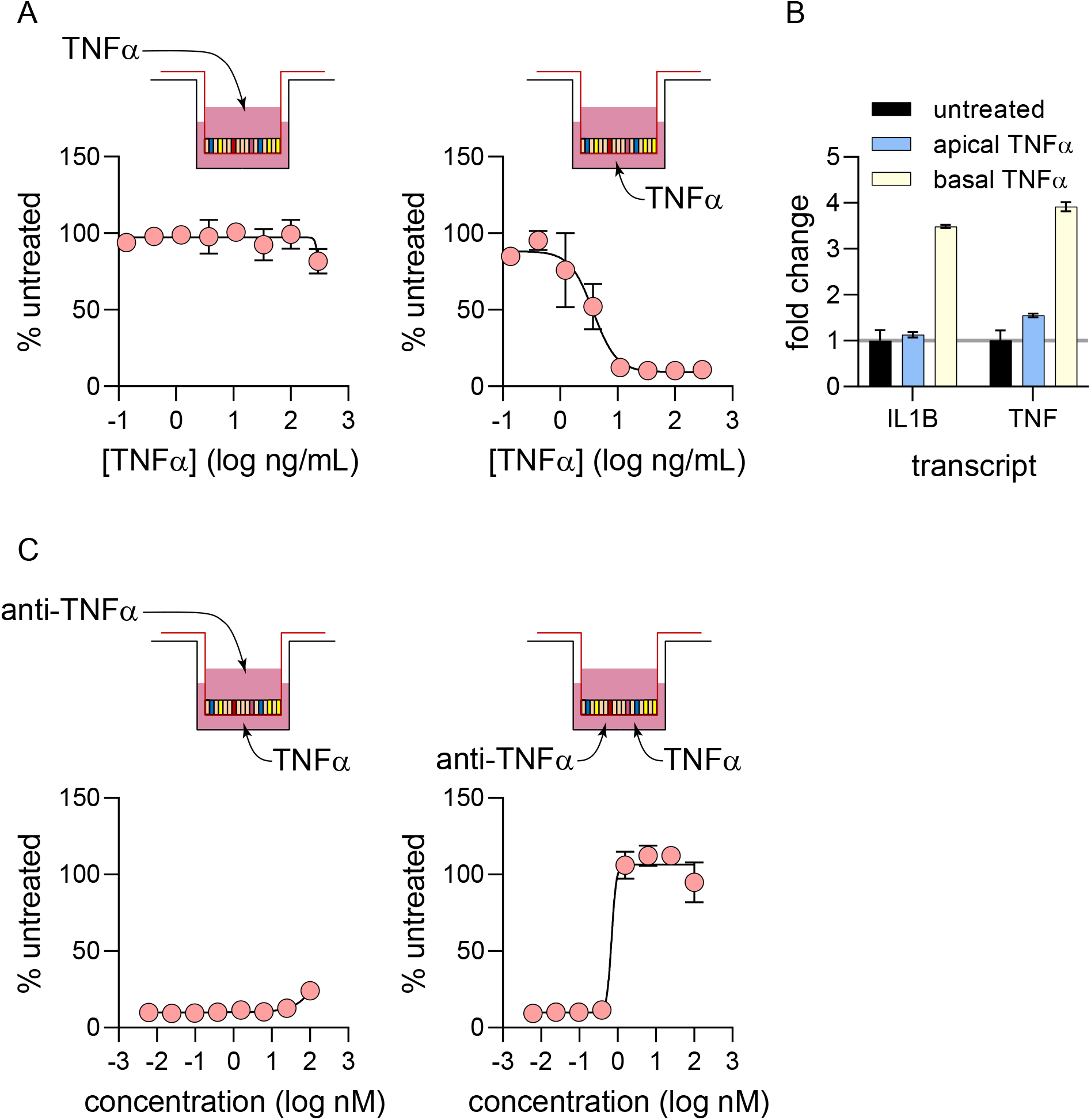
Apical vs basolateral TNF-α signaling (A) Monolayers were challenged with an 8-point dose curve of TNF-α on either the apical or basolateral sides of the monolayer. TEER measurements were taken at 72-hours post-challenge and analyzed in a similar manner to Figure 2B. (B) Quantifying differences in transcriptional response to 40 ng/mL TNF-α challenge on either side of monolayers. 6 hours post-challenge, monolayers were lysed and processed for qPCR. Gene expression for IL1B and TNF were quantified, and fold change values were calculated by normalizing to untreated controls. (C) Quantifying the potency of anti-TNF-α neutralizing antibodies by different routes of delivery. Monolayers were pre-treated with anti-TNF-α antibody on either apical or basolateral sides before TNF-α challenge on the basolateral side. TEER measurement was taken 72 hours post-challenge and EC50 values and curve fit was performed in the same manner as Figure 2C.

Next, we investigated the directionality of barrier function rescue by anti-TNF-α antibody. TNF-α was added to the basolateral chamber and anti-TNF-α antibody was added either in the apical or basolateral chamber. Addition of anti-TNF-α antibody to the apical chamber did little to rescue the barrier from damage by TNF-α (Figure 3C). Conversely, addition of anti-TNF-α to the basolateral chamber rescued barrier damage in a dose dependent manner (EC50 = 0.7 nM; Figure 3C), similar to results with antibody added to both chambers (EC50 = 1 nM; Figure 2C). This experiment showed that the vast majority of signal seems to have initiated from the basolateral compartment in this culture system.

### Quantifying receptor-proximal phosphorylation events in organoid-derived monolayer cultures

TNF-α-induced barrier damage is a distal consequence of proximal events such as receptor binding and changes in gene expression and cytokine/chemokine secretion patterns. Therefore, we sought to develop a clear understanding of early events that shape eventual disruption of barrier function in organoid-derived monolayers. As a receptor-proximal event, phosphorylation of p65 component of NFκB was quantified.

Upon receptor binding, TNF-α mediates NFκB pathway activation. A key molecular event in the activation of this pathway is the phosphorylation of p65 shortly after receptor binding. Phosphorylation is a labile event, so a time course experiment was designed to determine the peak phosphorylation for p65. TNF-α stimulation led to a sharp increase in p65 phosphorylation, peaking at 15 minutes (Figure 4A). After this peak, the signal slowly tapered off back to background levels by 120 minutes post-stimulation (Figure 4A). Next, organoid-derived monolayers were stimulated with multiple TNF-α concentrations showing a dose-dependent increase in signal (EC50 = 48 ng/mL; Figure 4B). Importantly, TNF-α-induced p65 phosphorylation could be inhibited by the addition of anti-TNF-α antibody (IC50 = 0.9 nM; Figure 4C). These results indicate that in organoid-derived monolayers, receptor engagement by TNF-α leads to NFκB pathway activation through p65 phosphorylation.

**Figure 4:**
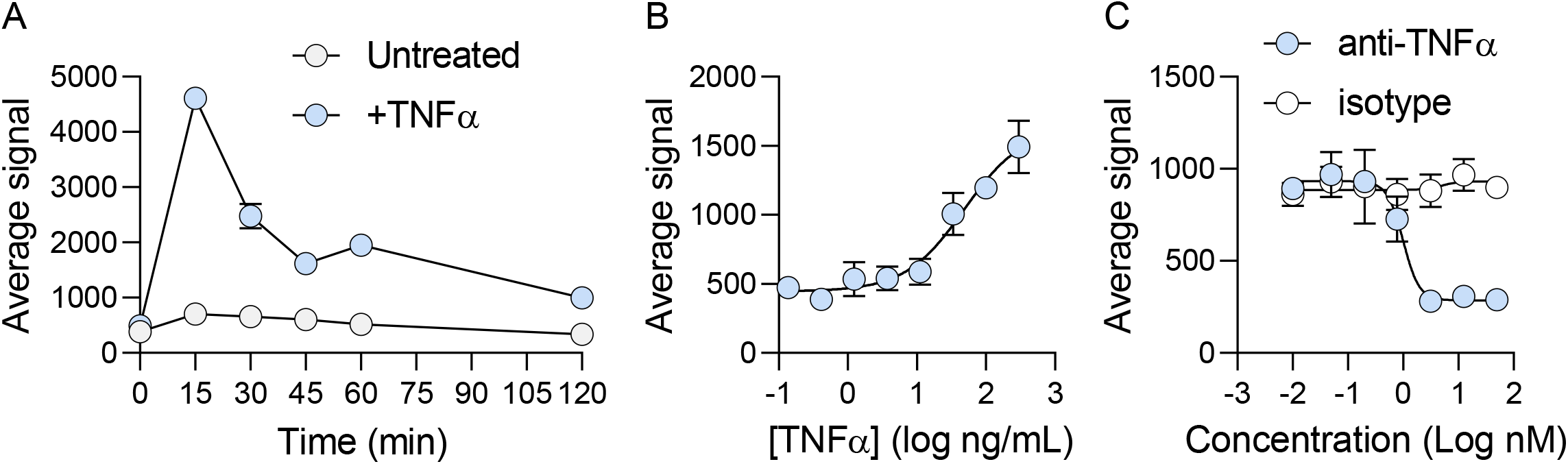
Quantifying p65 phosphorylation in response to TNF-α stimulation of organoid-derived monolayer cultures (A) Organoid-derived monolayers were left untreated or stimulated with 40 ng/mL TNF-α. At indicated time points, cells were lysed and levels of phosphorylated p65 was quantified using an MSD assay. (B) A dose-series of TNF-α was added to organoid-derived monolayers. At 15 min post stimulation cells were lysed and phospho-p65 was quantified. (C) Organoid-derived monolayers were incubated with either an anti-TNF-α antibody or an isotype-matched control for 1 hr. TNF-α (40 ng/mL) was added to all samples and cells were harvested by lysis after 15 min of stimulation, followed by phospho-p65 quantification.

To understand the extent by which organoid-derived monolayers can recapitulate inflammatory signaling, the same proximal target phosphorylation was performed in THP-1 cells, a known cellular model commonly used in IBD research due to their high reproducibility and ability to accurately simulate inflammation, notably for TNF-α signaling^27,28^. Consistently, we observed the strongest p65 phosphorylation at 15 minutes post TNF-α stimulation in THP-1 cells (Figure S1A). Importantly, anti-TNF-α antibody decreased p65 phosphorylation in a dose-dependent manner, similarly to the organoid-derived monolayers (IC50 = 0.158 nM; Figure S1B). Additionally, we confirmed the correlation of inhibiting p65 phosphorylation in organoid-derived monolayer cultures with MIP-1β inflammatory cytokine secretion, which is induced via TNFα-NF-κB signaling^29^, in the translational human whole blood assay (IC50 = 0.1514 nM, n = 11; Figure S1C). Altogether, these data show robust transcription factor phosphorylation in response to TNF-α.

### Target gene expression in response to TNF-*α* stimulation

The transcriptional response to TNF-α in organoid-derived monolayers was quantified next. Quantitative PCR was used to determine target gene expression profile. 2D organoid-derived monolayers were challenged with TNF-α for 6 hours followed by RNA purification and quantitative PCR. Using a TaqMan array for the NFκB pathway, the expression of 84 NFκB related genes was quantified. Of the genes in this array, 68 were expressed in the monolayer system (Figure 5A). Of these, 37 were upregulated significantly (p < 0.05) in the TNF-α-challenged cultures compared to untreated controls (Figure 5B). Many TNF-α response and NFκB signaling pathway components were significantly upregulated, including proximal components of the TNF-α response (TRAF2, NFKBIA), transcription factors in the Rel/NFKB family (RelB, NFKB1, NFKB2), and death-inducing signaling complex (FAS). Additionally, expression of pro-inflammatory cytokines that directly affect the intestinal epithelium, TNF-α (TNF) and IL-1β (IL1B), was significantly upregulated. Furthermore, expression for many cytokines and chemokines associated with immune cell recruitment or differentiation were significantly upregulated, including IL-8 (CXCL8), IP-10 (CXCL10), CD54 (ICAM1), CSF1, C3, CXCL2, CXCL1, CCL5, MDC (CCL22), and MCP1 (CCL2) (Figure 5B). These results indicate that TNF-α treatment led to robust transcriptional activation of NFκB signaling in organoid-derived monolayers.

**Figure 5:**
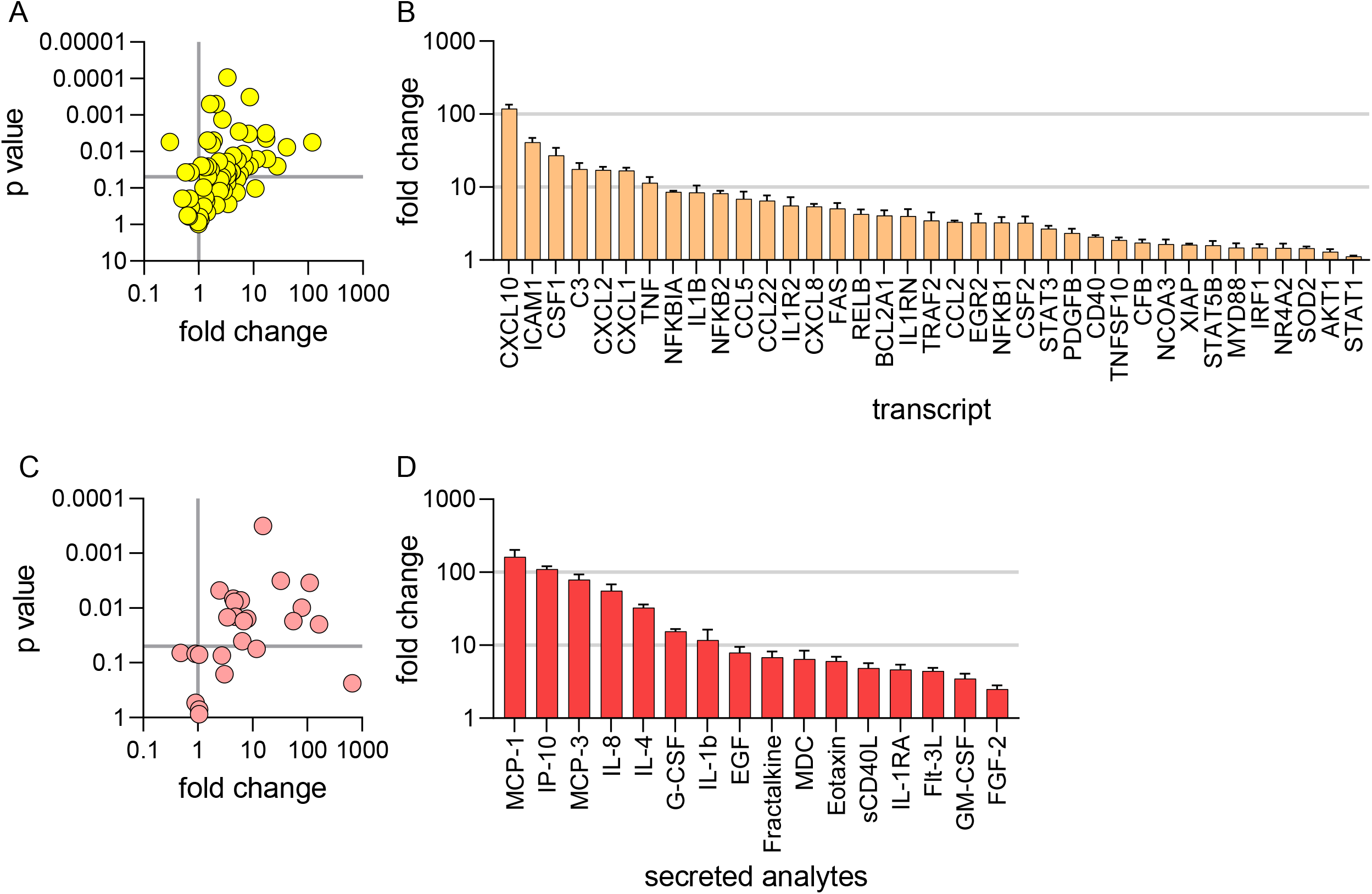
Target gene expression and cytokine production in response to TNF-α stimulation (A and B) Assessing changes in NFκB-related gene expression following TNF-α stimulation. Following 6-hour challenge with 40 ng/mL TNF-α on both apical and basolateral sides of the monolayer, a ratio of gene expression (by qPCR) compared to untreated controls was calculated for each marker in an 84-gene TaqMan array. Data were visualized as a volcano plot of p-value versus fold change (TNF-α-treated / untreated comparison), with each data point representing the average of 3 cell culture replicates (A). Markers with a fold change above 1 and p < 0.05 were determined to be significantly upregulated and ranked by fold change (B). (C and D) Assessing changes in the cytokine secretion profile of organoid-derived monolayers following TNF-α challenge. Monolayers were challenged with 40 ng/mL TNF-α on both apical and basolateral sides for 24 hours before supernatant collection and Luminex analysis. Similar to gene expression analysis, results were visualized as a volcano plot of p-value vs fold change. Each data point represents the average of 3 cell culture replicates (C). Markers with a fold change above 1 and p < 0.05 were determined to be significantly upregulated and ranked by fold change (D).

### Cytokine production in response to TNF-*α* stimulation

An experiment was performed to analyze the cytokine secretion profile of organoid-derived monolayers in response to TNF-α. Cultures were either untreated or challenged with TNF-α on the apical or basolateral side. Supernatants were collected from both apical and basolateral sides at 0, 3, 6, and 24 hours after stimulation and analyzed using a 38-plex human cytokine/chemokine Luminex panel. Of the 38 analytes included, 28 were detected in monolayer cultures at 24 hours (Figure 5C). 16 cytokines or chemokines were significantly upregulated at 24 hours in the supernatants that were collected in the basolateral side of the epithelium (Figure 5D). The supernatants collected from the apical chamber were also evaluated but showed much lower or undetectable levels of almost all analytes. Supernatant collection at 4 time points allowed for the generation of time course of cytokine secretion for these analytes. In agreement with the NFκB gene expression panel, many cytokines and chemokines associated with the recruitment and differentiation of immune cells were secreted in response to TNF-α, including MCP-1, IP-10, MCP-3, IL-8, IL-4, G-CSF, GM-CSF, MDC, fractalkine, and eotaxin. These results showed that stimulation of organoid-derived monolayers with TNF-α produced a consistent secreted analyte signature. We next determined whether cytokine/chemokine secretion could be employed to evaluate TNF-α pathway modulators.

### Assessing TNF-*α* pathway modulators using IL-8 secretion

We showed that in response to TNF-α, organoid-derived monolayer cultures secrete a variety of cytokines and chemokines, modelling the inflammatory cascade that occurs in IBD. Notably, IL-8 was shown to be consistently secreted into the basolateral chamber in response to TNF-α stimulation (Figure 5D). IL-8 is a key factor in amplifying the inflammatory response in IBD as it is a potent chemoattractant of neutrophils ^30^. The observed production of IL-8 in response to TNF-α stimulation is expected and further validates the physiological relevance of this organoid-derived monolayer system. Next, we sought to further characterize the IL-8 response to TNF-α stimulation with the goal of quantifying the effects of TNF-α pathway modulators upon IL-8 production.

First, organoid-derived monolayers were challenged with a dose curve of TNF-α and supernatants were collected from the basolateral chamber at 24 hours and analyzed using an IL-8 ELISA. A dose-dependent increase in IL-8 secretion was observed (Figure 6A), similar to the response observed in the TEER assay (Figure 2B). EC50 values for the two experiments were similar (7 ng/mL for TEER and 11 ng/mL for IL-8 secretion) indicating concordance between assays.

**Figure 6:**
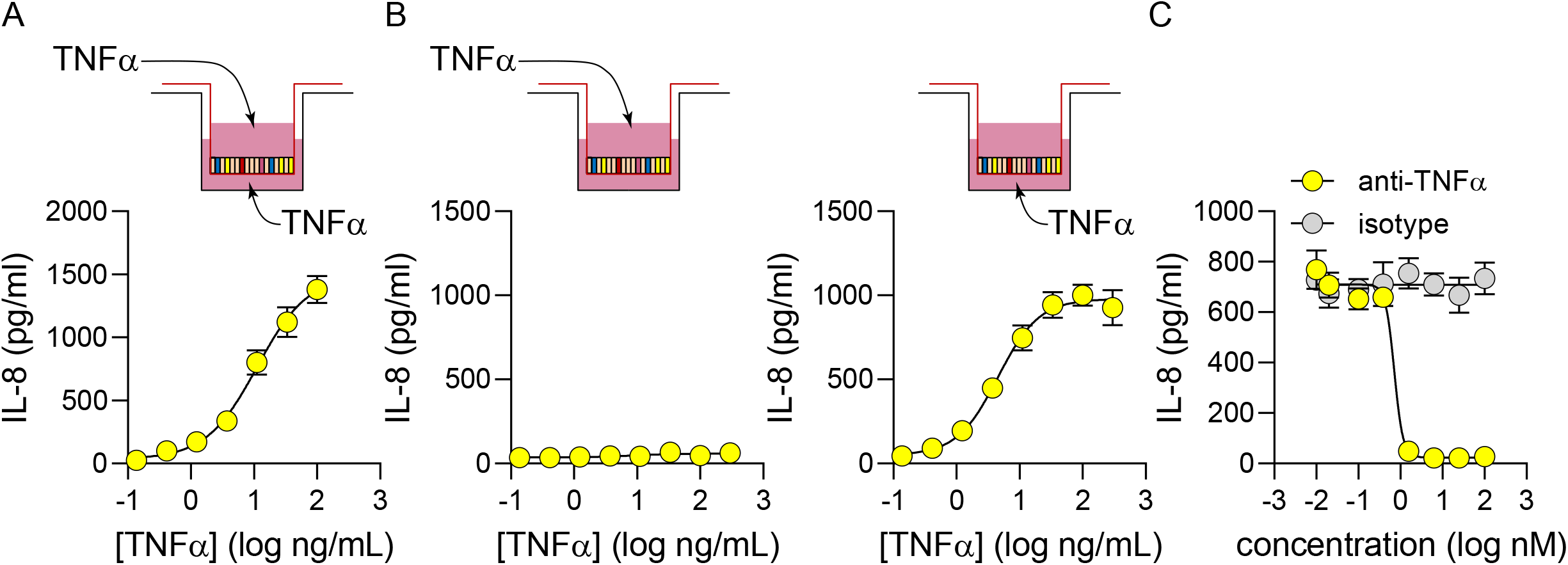
Assessing TNF-α pathway modulators by quantifying IL-8 secretion. Secretion of IL-8 into basolateral chamber was quantified after 24 hours challenge with a dose range of TNF-α on both apical and basolateral surfaces of organoid-derived monolayers. Secretion of IL-8 into basolateral chamber was quantified 24 hrs after stimulation on either apical or basolateral surface. (C) Organoid-derived monolayers were pre-treated for 1 hr with either an anti-TNF-α antibody or isotype-matched control, then challenged with 40 ng/mL TNF-α on both apical and basolateral surfaces. IL-8 secretion was quantified in the basolateral chamber after 24 hrs. IL-8 amounts were quantified by ELISA, and each data point represents the average of 5 cell culture replicates. All curve fitting and potency values were generated using the same nonlinear regression model for the TEER measurements (Figures 2 and 3).

Next, the polarity of IL-8 production in response to TNF-α was determined. Monolayers were challenged with TNF-α from either the apical or basolateral side and IL-8 secretion quantified in the basolateral chamber. A dose-dependent IL-8 response was observed only upon TNF-α stimulation from the basolateral side of the epithelium (EC50 = 4.5 ng/mL; Figure 6B). Once again, this readout recapitulates TEER results (Figure 3A), as there was almost no signal observed with apical challenge with TNF-α. These results show that IL-8 secretion is a reliable indicator for TNF-α pathway activity, so the effect of pathway inhibitors was quantified next.

An anti-TNF-α antibody decreased IL-8 secretion levels in in TNF-α-stimulated monolayer cultures a dose-dependent manner (IC50 = 0.7 nM; Figure 6C). This half maximal value was once again similar to results obtained with the TEER assay (EC50 = 1 nM; Figure 2C). Taken together, these results indicate that IL-8 secretion is a dependable method to quantitatively assess the effect of TNF-α pathway modulators.

## Discussion

In this work we show the development of a high-throughput primary human culture system that recapitulates key features of intestinal inflammation. We demonstrate that organoid-derived monolayer cultures are amenable to quantitative assessment through multiple readouts while retaining important physiologically-relevant features (e.g. cell composition). We first adapted organoids to a monolayer cultures system and demonstrated efficient differentiation and barrier function (Figure 1). We showed that TNF-α reduced barrier function in a dose-dependent manner and that loss of barrier function could be rescued by the addition of TNF-α-pathway neutralizing antibodies (Figure 2). As epithelial cells are polarized in the human intestine, unidirectionality of TNF-α signaling was investigated. TNF-α-mediated barrier damage could be initiated only from the basolateral compartment, demonstrating that epithelial cell polarization was preserved in our organoid-derived monolayers (Figure 3). Additionally, we showed that key receptor-proximal signaling events (phosphorylation, target gene expression, cytokine/chemokine production) were intact and that these events could be used as quantitative readouts for receptor engagement (Figures 4-6). Taken together, we have demonstrated that organoid-derived monolayer cultures are a tractable system to investigate gut inflammatory indications. Importantly, we have established molecular and phenotypic readouts in this system that span a large time scale after receptor engagement, from minutes (phosphorylation) to hours (target gene expression, cytokine/chemokine production) to days (barrier dysfunction).

Importantly, in addition to miniaturization and automated readouts, this platform reaches a mature (differentiated) state in one week. Most Caco-2 systems take considerably longer to mature after plating, and still do not recapitulate many key aspects of intestinal epithelial biology. Conversely, in addition to more rapid maturation, this organoid-derived platform can generate multiple functional cell types found in the intestinal epithelium, develop a physiological barrier function, and appropriately responds to inflammatory stimuli such as TNF-α. A number of organoid-derived monolayer systems have been described recently ^17-21,31-38^.

These systems highlight the vast potential of primary epithelial cell systems to recapitulate various aspects of intestinal biology and disease. These range from investigating transporter function to pharmacokinetic studies, to co-culture of epithelial cells with pathogens and/or immune/stromal cells. While similar to the system described in our paper, previously published publications have not focused on teasing apart inflammatory signaling pathways using multiple readouts in a 96-well format. Nonetheless, previous publications point to potential future advancement opportunities for our monolayer system. Notably, our system is composed solely of epithelial cells. An important feature of intestinal inflammation is the cross-cell type communication, mediated by immune and stromal cells, which is an important feature of initiating and maintaining an inflamed state. Incorporation of immune and/or stromal cells into our system will steer the model even closer to recapitulating important aspects of human disease.

This miniaturized 96-well scale, combined with standard automation, allows for higher throughput screening that is not commonly associated with complex primary cell system. This organoid-derived monolayer system is physiologically-relevant, yet practical. All reagents, consumables, and equipment used are commercially available. This is a substantial advantage over most organ-on-chip platforms that require specialized fabrication, complicated laboratory setups, and extensive expertise and effort to implement. This system was designed to recapitulate cell composition and key inflammatory responses of the human intestine in a format that is amenable to robust hypothesis generation and testing in a quantitative manner.

## Materials and Methods

### ASC organoid culture and monolayer seeding

Adult stem cell (ASC) intestinal organoid cultures were established by isolating crypts from human intestinal tissue as previously described ^22-24^. These ASC organoids were maintained and expanded by passaging once per week in a proliferation medium (50% L-WRN conditioned media + Y27632 + SB431542 ^24^). 2D organoid-derived monolayers were generated from these 3D cultures. On day 5 post-passaging, 3D organoids were collected in serum-free washing media (Advanced DMEM/F12; Invitrogen 12634) and removed from Matrigel by repeated pipetting.

Organoids were broken down completely into single cells by suspending in TrypLE Express (Gibco 12605010) and incubating at 37ºC for 15 minutes, followed by repeated pipetting. After washing to remove TrypLE, single cells were suspended in proliferation media and seeded at a density of 0.05E6 cells/well for 96-well transwells (MilliporeSigma PSHT004R5), 0.15E6 cells/well for 24-well transwells (Corning 3460), or 0.45E6 cells/well for 12-well transwells (Corning 3470). Before seeding, transwells were coated with 0.35 mg/mL Matrigel Membrane Matrix (Corning 354234) diluted with DPBS (Gibco 14190144) for at least 1 hour at 37C. After coating time was complete, transwells were washed once with DPBS before plating the cells.

12-well, 24-well, and 96-well experiments were all used throughout various stages in this paper. The 12-well transwell system was used for all imaging experiments, 24-well transwell system was used for gene expression and cytokine secretion analysis, and 96-well transwell system was used for phospho-p65 quantification, cytokine secretion, and all trans-epithelial electrical resistance (TEER) measurement analyses. In all formats, 0.4 um pore sizes were used, and the membrane material was either polyester (12- and 24-well) or polycarbonate (96-well). After seeding, organoid-derived monolayers were grown in proliferation media for 4 days before switching to differentiation media (5% L-WRN conditioned). Monolayers were then grown in differentiation media for 4 days before typically initiating experiments on day 8 after seeding. Media was changed every 1-2 days.

### TNF-*α*-mediated barrier damage assay

To assess TNF-α-mediated barrier damage, fully differentiated monolayers were challenged with a dose curve of TNF-α (R&D Systems 210-TA-100). Anti-TNF antibody (Invivogen htnfa-mab1) was evaluated by pre-treating for 1 hour before TNF-α challenge. TEER measurements occurred immediately after challenge and in 24-hour intervals thereafter, with the 72-hour timepoint being the final and displayed data. TEER measurements were performed using the REMS Autosampler produced by World Precision Instruments. Raw resistance values generated by the system were corrected to TEER values by normalizing to transwell surface area, and values for challenged conditions were further normalized to untreated samples to generate “% untreated” values. Curve fitting and EC50 values were generated in GraphPad Prism using a 4-parameter nonlinear regression model.

### Transcriptional analysis

To evaluate organoid-derived monolayer growth and differentiation, cultures were harvested for RNA purification and qPCR analysis at proliferative and fully differentiated timepoints (days 3 and 8 after plating, respectively). To evaluate gene expression changes in response to TNF-α, fully differentiated cultures were challenged for 6 hours before harvesting for RNA purification and qPCR analysis. In both cases, Ct values were normalized to endogenous controls and presented as fold change of differentiated/challenged cultures over proliferative/untreated controls. TaqMan assays used in this study are listed in supplemental table 1. The “Human NFκB Signaling” TaqMan array (Invitrogen 4413255, design ID: RPGZE7K) was used to quantify expression of NFκB target genes.

### Phosphorylation assay

Cultures were challenged with TNF-α with or without pre-treatment of an anti-TNF-α antibody. At indicated timepoints, cells were lysed (lysis buffer: Cell Signaling Technology 9803) and frozen. Quantification of NFκB p65 phosphorylation was performed using a Phospho-p65 (Ser536) Whole Cell Lysate Kit (Meso Scale Diagnostics K151ECD) according to the manufacturer protocol.

### Immunofluorescence microscopy

For the detection of markers of functional cell types or junction proteins, fully differentiated organoid-derived monolayer cultures were fixed using 4% paraformaldehyde, permeabilized with 1% Triton X-100, and blocked with 5% BSA. Monolayers were then incubated overnight at 4°C with primary antibodies for markers of interest, followed by secondary antibody staining at room temperature for 2 hours. Stained monolayers were mounted onto slides and microscopy was performed with a confocal microscope. To stain for proliferative cells, an EdU assay was performed according to the manufacturer’s protocol. All reagents are listed in supplemental table 2.

### Cytokine secretion analysis

Supernatants were collected from samples from the basolateral side of transwells at indicated timepoints after TNF-α challenge. Cytokine levels in supernatants were analyzed by 38 plex MILLIPLEX MAP Human Cytokine/Chemokine Magnetic Bead Panel (Millipore HCYTMAG-60K-PX38) adapted for use with 96-well DropArray plates and DropArray LT210 washing station (Curiox Biosystems). IL-8 secretion levels were quantified using an IL-8 ELISA kit (R&D Systems D8000C). Curve fitting and potency values for IL-8 secretion data were generated using a 4-parameter nonlinear regression model in GraphPad Prism.

## Acknowledgements

We thank Drs. Stephen Kasper, Katharine Best, and Marc Levesque for critical reading of the manuscript.

## Author Contributions

C. W. W. designed experiments, acquired, analyzed, and interpreted results, and drafted the manuscript; N. L., acquired and analyzed data for Figures 4 and S1 and revised the manuscript; L. S., and A. H. acquired and analyzed data; N. B., S. A., S. V., and K. B., and L. A. L. interpreted the results and revised the manuscript; S. M. conceived and designed the work, acquired, analyzed, and interpreted results, and wrote and revised the manuscript.

## Data Availability Statement

No ‘omics’ datasets were generated. The datasets generated in the current study are available from the corresponding author on reasonable request.

## Figure legends

**Supplemental Table 1:** TaqMan probes used in this study

| transcript | TaqMan ID | marker type |
| --- | --- | --- |
| PPIB | Hs00168719_m1 | endogenous control |
| MKI67 | Hs01032443_m1 | proliferative/stem cells |
| LGR5 | Hs00969422_m1 |  |
| AXIN2 | Hs00610344_m1 |  |
| MUC2 | Hs03005103_g1 | secretory cells |
| TFF3 | Hs00902278_m1 |  |
| CHGA | Hs00900370_m1 |  |
| MMP7 | Hs01042796_m1 |  |
| P-gp/ABCB1/CD243 | Hs00184500_m1 | absorptive cells |
| BCRP/ABCG2 | Hs01053790_m1 |  |
| SLC15A1 | Hs00192639_m1 |  |
| SLC10A2/ASBT | Hs01001557_m1 |  |
| TNFR1/TNFRSF1A | Hs01042313_m1 | immune receptors |
| TNFR2/TNFRSF1B | Hs00961750_m1 |  |
| IL1B | Hs01555410_m1 | cytokines |
| TNF | Hs00174128_m1 |  |

**Supplemental Table 2:** Fluorescence microscopy reagents

| Reagent | Dilution | Source | Catalog # |
| --- | --- | --- | --- |
| Mouse anti-human Mucin 2 antibody | 1:200 | Dako | M731329-2 |
| Rabbit anti-human Lysozyme antibody | 1:500 | Invitrogen | PA5-16668 |
| Rabbit anti-human Chromogranin A antibody | 1:100 | Abcam | AB15160 |
| Hoechst 33342 | 1:2000 | Invitrogen | H3570 |
| Goat anti-rabbit Alexa 488 | 1:250 | Invitrogen | A-11034 |
| Goat anti-mouse Alexa 594 | 1:250 | Invitrogen | A-11032 |
| Click-iT EdU Cell Proliferation Kit for Imaging, Alexa Fluor 594 | N/A | Invitrogen | C10339 |

**Figure S1:**
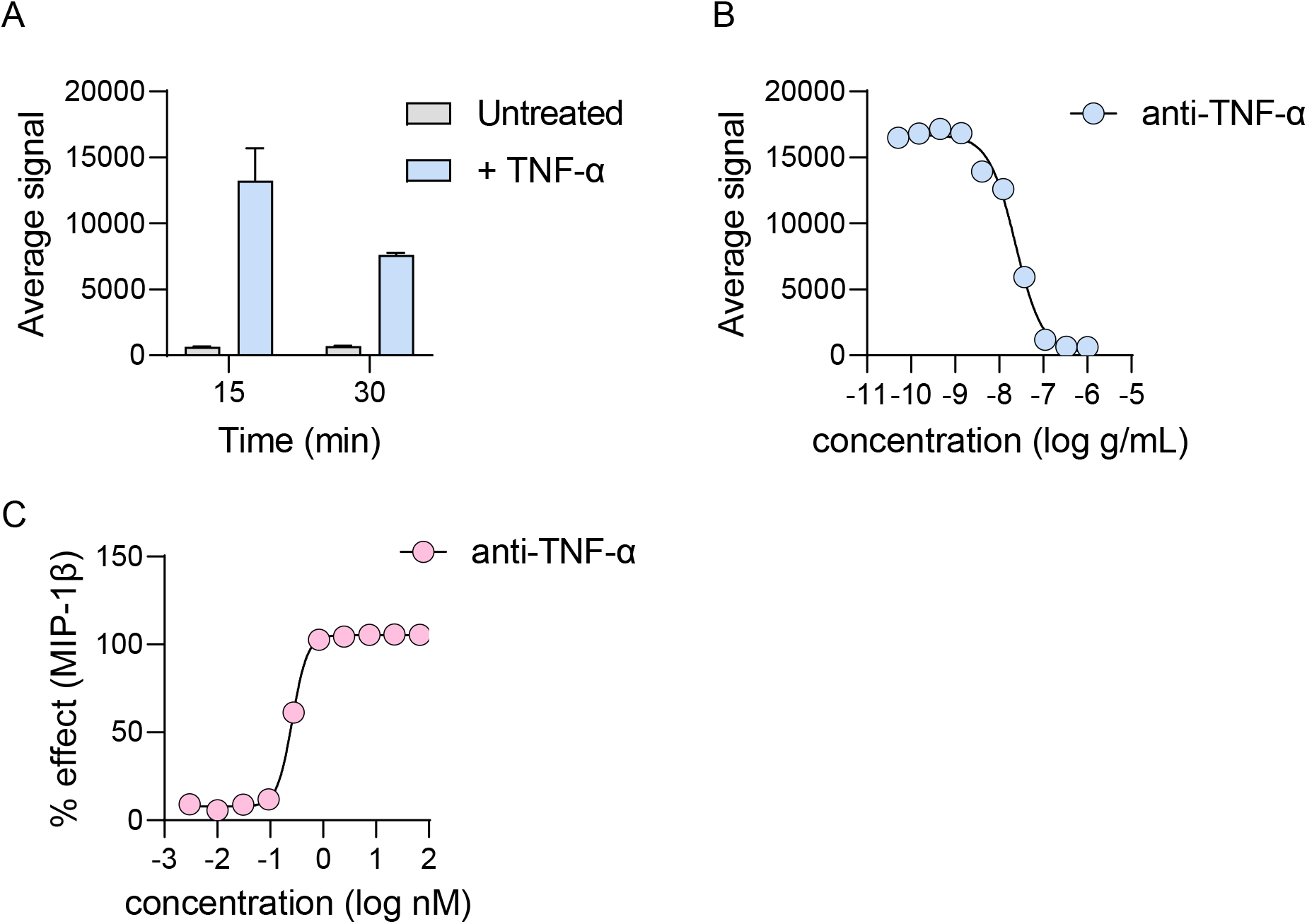
Quantifying p65 phosphorylation and MIP-1β secretion in THP-1 and human whole blood (A) THP-1 cells were untreated or treated with 10 ng/mL of TNF-α. Cells were lysed at the indicated timepoints, and phosphorylated p65 was quantified by MSD assay. (B) THP-1 cells were treated with a dose range of anti-TNF-α in the presence of 10 ng/mL of TNF-α. Phosphorylated p65 was quantified by MSD assay. (C) Human whole blood was treated with 10 ng/mL of TNF-α and dose-series of anti-TNF-α antibody. MIP-1β secretion was quantified by MSD assay.

